# Host response of Syrian hamster to SARS-CoV-2 infection, including differences with humans and between sexes

**DOI:** 10.1101/2022.11.22.517339

**Authors:** Martina Castellan, Gianpiero Zamperin, Giulia Franzoni, Greta Foiani, Maira Zorzan, Petra Drzewnioková, Marzia Mancin, Irene Brian, Alessio Bortolami, Matteo Pagliari, Annalisa Oggiano, Marta Vascellari, Valentina Panzarin, Sergio Crovella, Isabella Monne, Calogero Terregino, Paola De Benedictis, Stefania Leopardi

**Affiliations:** Division of Comparative Biomedical Sciences, Istituto Zooprofilattico Sperimentale delle Venezie, Legnaro, Italy; Laboratory for Emerging Viral Zoonoses, Istituto Zooprofilattico Sperimentale delle Venezie, Legnaro, Italy; Viral genomics and transcriptomics Laboratory, Istituto Zooprofilattico Sperimentale delle Venezie, Legnaro, Italy; Laboratory of Diagnostic Virology, Istituto Zooprofilattico Sperimentale della Sardegna; Laboratory of Histopathology, Istituto Zooprofilattico Sperimentale delle Venezie, Legnaro, Italy; Risk Analysis and Public Health Department, Istituto Zooprofilattico Sperimentale delle Venezie, Legnaro, Italy; Innovative Virology Laboratory, Istituto Zooprofilattico Sperimentale delle Venezie, Legnaro, Italy; Laboratory of Experimental Animal Models, Istituto Zooprofilattico Sperimentale delle Venezie, Legnaro, Italy; Biological Science Program, Department of Biological and Environmental Sciences, College of Arts and Sciences, Qatar University, Doha

**Keywords:** SARS-CoV-2, animal model, sex, histopathology, host response, immune response, transcriptomic profile

## Abstract

The emergence of severe acute respiratory syndrome coronavirus 2 (SARS-CoV-2) has highlighted the importance of having proper tools and models to study the pathophysiology of emerging infectious diseases to test therapeutic protocols, assess changes in viral phenotype and evaluate the effect of viral evolution. This study provides a comprehensive characterization of the Syrian hamster (*Mesocricetus auratus*) as an animal model for SARS-CoV-2 infection, using different approaches (description of clinical signs, viral load, receptor profiling and host immune response) and targeting four different organs (lungs, intestine, brain and PBMCs). Our data showed that both male and female hamsters are susceptible to the infection and develop a disease similar to the one observed in patients with COVID-19, including moderate to severe pulmonary lesions, inflammation and recruitment of the immune system in lungs and at systemic level. However, all animals recovered within 14 days without developing the severe pathology seen in humans, and none of them died. We found faint evidence for intestinal and neurological tropism associated with the absence of lesions and a minimal host response in intestines and brains, highlighting another crucial difference with the multi-organ impairment of severe COVID-19. When comparing male and female hamsters, it was observed that males sustained higher viral RNA shedding and replication in the lungs, suffered from more severe symptoms and histopathological lesions and triggered higher pulmonary inflammation. Overall, these data confirm the Syrian hamster as a suitable model for mild-moderate COVID-19 and reflect sex-related differences in the response against the virus observed in humans.

## 1. Introduction

The pandemic of coronavirus disease 2019 (COVID-19) has resulted in a devastating global threat to human society, economy and healthcare system [1–3]. The disease is caused by severe acute respiratory syndrome coronavirus 2 (SARS-CoV-2), a positive-sense single-stranded RNA virus belonging to the subgenus *Sarbecovirus*, genus *Betacoronavirus*, species *SARS-related coronavirus*, emerged from a animal source after zoonotic cross-species transmission [4–6]. The virus mostly replicates in the respiratory tract, although patients may also experience disorders associated to multi-organ engagement, including neurologic and gastro enteric symptoms, whose incidence, mechanism and significance are still a matter of discussion [7–11]. According to the age of the patient and the presence of predisposing factors, COVID-19 varies widely in the severity of its clinical manifestations, spanning from asymptomatic infections to an acute respiratory distress syndrome (ARDS) requiring mechanical ventilation and, in the worst-case scenario, leading to death [12–14]. Epidemiological data indicate that males are more prone to develop a severe COVID-19 symptomatology, suggesting that sex may also influence SARS-CoV-2 pathogenesis due to genetic and hormonal factors, although social-behavioral differences between genders may also play a role [15–18]. Regardless of the cause, the severe form of the disease follows a common mechanism in the dysregulation of the inflammatory response, similarly to what has been observed with other coronavirus infections, such as Severe Acute Respiratory Syndrome (SARS) and Middle East Respiratory Syndrome (MERS) [12]. Briefly, in the attempt to clear the infection, the immune system of certain individuals releases an excessive amount of pro-inflammatory cytokines, known as “cytokine storm”, promoting an uncontrolled inflammation that damages lungs and other organs, such as brain, gut and heart [19,20]. This important evidence has paved the way for a diagnostic, prognostic and therapeutic approach focused on controlling patients’ immune response, with particular attention to the innate immunity [20,21]. The overarching goal is to control the pandemic by reducing the incidence of severe manifestations through vaccination campaigns, and to develop and assess the efficacy of therapeutic agents against the new variants deriving from the evolution of SARS-CoV-2. Among them, the WHO classifies as “variants of concern (VOCs)” viruses [22] that show mutations on the spike protein that might influence transmissibility, symptomatology, immune-evasion, efficacy of therapeutic monoclonal antibodies and sensitivity of diagnostic methods [23– 30]. The first variant defined as VOC was the Alpha, also referred to as B.1.1.7, which, compared to older strains, was associated with higher transmissibility and, according to some studies, increased mortality rates [31]. The Alpha VOC was first described in the United Kingdom in November 2020 and rapidly spread across Europe, being responsible for an increased number of infections during the second epidemic wave. In Italy, it was the most prevalent variant between February and March 2021 [32,33].

In the race against a pandemic event, researchers need reliable animal models that i) are susceptible to the infection, ii) are able to eliminate the virus, iii) display clinical and pathological manifestations typical of human disease, and iv) mimic the same immune impairments found in patients. Non-human primates, ferrets, hamsters, and transgenic mice (i.e. K18-hACE2 mouse) are permissive to SARS-CoV-2 infection and develop lung lesions resembling the pathological patterns found in humans [31,34–39]. Among these, the Syrian hamster (*Mesocricetus auratus*) exhibits the best balance between costs, neurological development, availability, easy handling and maintenance in captivity and it is extensively used for translational medicine [31,36,40,41]. Previous studies showed that SARS-CoV-2 replicates efficiently in the respiratory tract of hamsters and is able to invade the central nervous system, with no differences observed between animals of different age [40,42]. Histopathological and radiographic evaluations confirmed that these animals develop pneumonia without showing severe clinical manifestations and fully recover in 2-3 weeks, albeit chronic sequelae can be seen in the lungs, kidney and the olfactory bulb up to 31 days post infection [40,42,43]. In addition, preliminary studies showed that hamsters increase the gene expression of some cytokines/chemokines in the lungs, which may be compatible with the cytokine storm described in humans [42,44]. The aim of this study is to provide an in-depth evaluation of the Syrian hamster as animal model for human COVID-19, and to identify the advantages and disadvantages of using this species for translational medicine. Our work provides new outcomes that need to be taken into account while designing an infection study using this animal model, including evidence for sex-related peculiarities and critical differences with the human host response.

## 2. Materials and Methods

### 2.1. Animal experiment

The study involved 60 8 weeks old Syrian hamsters, divided into 4 experimental groups of 15 individuals, each composed of only females or only males, either infected or used as negative controls. Animals were acclimatized 7 days prior to infection in individual cages (BCU-2 Rat Sealed Negative Pressure IVC, Allentown Inc) in the biosafety level 3 (BSL3) facility, following national and international regulations on the welfare of laboratory animals.

Animals of the infected groups were inoculated intranasally under general anesthesia with Isoflurane using SARS-CoV-2 B.1.1.7 (Alpha) variant (Accession N: EPI_ISL_766579), isolated in 2020 from a mild/severe COVID case as previously described [45]. The viral dose was of 8×10^4^ PFU/100µl, within the range used in the literature and known to induce productive infection in this model[40,43,46]. Hamsters included in the two groups of negative controls were inoculated using 100μl of sterile PBS solution.

Animals were daily monitored for 14 days to record clinical signs and general signs of distress, thus securing animal welfare besides collecting research data. At 2, 4, 6, 9 and 14 dpi we registered weights and collected oropharyngeal swabs and blood samples from the gingival vein under general anesthesia [47]. On day 2, 6 and 14 we euthanized 5 individuals per group and performed an intra-cardiac terminal blood collection for PBMCs isolation with Ficoll-Paque Plus (GE healthcare). We performed a complete necropsy of all animals and collected samples of the lungs, brains and intestines, using the best practices to avoid cross-contamination between different organs and animals. Specimens were fixed in 10% neutral-buffered formalin and in RNA later (Thermo Fisher®) for histological examination and molecular analyses respectively (Figure S3). Further details on tissue collection can be found as supplementary materials.

### 2.2. Histology and immunofluorescence

Formalin fixed samples were paraffin-embedded, cut in 4µm-thick sections and stained with hematoxylin and eosin (H&E) to evaluate the presence and severity of lesions in different organs. Histopathological lesions were scored as described elsewhere [44] (Table S2); total histopathological score has been defined as the sum of the scores attributed to each specific lesion. Slides were analyzed and images were taken using a Leica DM4 B light microscope with a DFC450 C Microscope Digital Camera at 20X and the software Leica Application Suite V4.13 (Leica Microsystems, Wetzlar, Germany).

We investigated the presence of the virus within tissues by immunofluorescence, using anti-SARS-CoV-2 spike glycoprotein and anti-dsRNA [48,49] as primary antibodies, in order to discriminate the presence of the antigen by the active replication of the virus within tissues, based on the fact that dsRNA is widely known as replicative intermediate for coronaviruses [50]. Immunofluorescence was also applied to investigate the expression of the ACE2 receptors within the tissues. Further details can be found as supplementary materials.

### 2.3. Molecular analyses for SARS-CoV-2 detection and quantification

The presence of viral RNA in oropharyngeal swabs was detected in all the control and infected individuals by qualitative rRT-PCR using the AgPath-ID™ One-Step RTPCR Reagents (Life Technologies) on a CFX96 Touch Deep Well Real-time PCR Detection System (Biorad). To quantify SARS-CoV-2 in target organs, we developed a digital droplet RT-PCR (RT-ddPCR) employing the One-Step RT-ddPCR Advanced Kit for Probes (Bio-Rad) and the QX200 Droplet Digital PCR System (Biorad). This approach was implemented only for the three individuals per experimental group that were randomly selected for transcriptomic analyses. Quantitative data were expressed as Log2 copies/ml of genomic RNA. Both tests targeted SARS-CoV-2 envelope protein (E) gene [51]. The quality of the samples was verified by amplification of the β-actin mRNA [52]. Further details can be found as supplementary materials.

### 2.4. Gene expression analyses by RNA-Seq

We randomly selected three individuals among five of each experimental group to investigate virus-host response by performing the transcriptomic profile of the lungs, brains, intestines and PBMCs of infected *versus* mock animals at three different time points along the infection, representing early infection, the infection apex and recovery.

Libraries were prepared with the Truseq Stranded mRNA library preparation kit (Illumina), following the manufacturer’s instructions and were run on an Agilent 2100 Bioanalyzer using an Agilent High Sensitivity DNA kit (Agilent Technologies) to ensure the proper range of cDNA length distribution. Sequencing was performed on Illumina NextSeq with NextSeq® 500/550 High Output Kit v2.5 (300 cycles; Illumina) in pair-end [PE] read mode, producing about 33 million reads per sample. After filtering raw data, we aligned high-quality reads against the reference genome of *Mesocricetus auratus* (BCM Maur 2.0, NCBI) [53] using STAR v2.7.9a [54] and generated the gene count using htseqcount v0.11.0 [55]. We then investigated the differential expression of genes between infected and mock males and females at each time point with Deseq2 package [56] and assigned Gene Ontology (GO) terms to each gene using Blast2GO v5.2.5 [57]. Child-father relationships belonging to GO graph were reconstructed using the OBO file downloaded from http://geneontology.org/ (accessed on 19/10/2021). Orthologs with *Homo sapiens, Mus musculus* and *Rattus norvegicus* were computed using Orthofinder v2.5.4 [58] and their proteome downloaded from Ensembl. Further details can be found as supplementary materials.

### 2.5. Serological analyses

In order to evaluate sero-conversion dynamics, we performed the focus reduction neutralization test (FRNT) as previously described, using for the detection of neutralizing antibodies the same viral strain used for the infection [59,60]. We defined as serum neutralization titer the reciprocal of the highest dilution resulting in a reduction of the control focus count higher than 90% (FRNT90). Sera of all animals were collected at 2, 4, 6, 9 and 14 dpi; only 4 out of 5 sera were collected for both males and females at 9 dpi.

We further analyzed serum samples of all the controls and infected animals at 2, 4, 6, 9 and 14 dpi for the presence of pro-inflammatory cytokines. Levels of IL-1β and IL-6 were assessed at the Istituto Zooprofilattico Sperimentale della Sardegna through singleplex ELISA using target-specific ELISA kits (MyBiosource), according to the manufacturers’ instructions and using an Epoch microplate reader (BioTek) to read absorbance.

### 2.6. Statistical analyses

We adopted the minimum sample size that guaranteed effective comparison between groups while minimizing the use of experimental animals. Infection of 13 out of 15 individuals per group indicated successful infection with a first type error α=0.01 (one tail) and a power 1-β=0.85.

To describe the shedding of viral RNA in males and females and to record changes of weights in hamsters during infection of males and females *versus* the corresponding group of mock animals, we performed descriptive statistical analysis using SAS 9.4 software [61–63]. We applied a spline mixed model by sex, taking into account the correlation between different observations of the same hamster over time using a first-order autoregressive AR(1) structure for the covariance matrix. We applied type III F-test to evaluate the significance of the overall effect of fixed factors specified in the model and performed post-hoc pairwise comparisons for each fixed factor of the mixed models to clarify differences. We performed studentized residuals analysis to assess the quality of the model and plotted QQ-plot, residuals distribution and scatterplot of the residuals versus the fitted to verify the assumption of normality and homoscedasticity. P-values of < 0.05 were considered significant, including Sidak adjusted P-values provided for multiple tests. For all the remaining statistical analyses, we used the Wilcoxon-Mann-Whitney test for independent groups implemented in GraphPad Prism 9. These included comparison i) between median antibody titers and serum levels of IL-1β and IL-6 of females *versus* males at each time point, ii) of histopathological total scores between infected *versus* mock animals of each sex and between infected females *vs* infected males at 2, 6 and 14 dpi, iii) of intra-alveolar inflammatory cell infiltration and perivascular/alveolar edema histopathological scores (lesions more abundant in males, see Results section) between infected *versus* mock animals of each sex and between infected females *vs* infected males at 2, 6 and 14 dpi. For all statistics, we considered as significant P*-*values < 0.05.

### 2.7. Data availability

RNA-Seq raw data generated in the present study were deposited in SRA under accession number PRJNA839918. Source data for Figures 1a-b; 2d-f; 3a-c; 4a-c; 5a–b; 6a, c are provided as Table S1-5.

**Figure 1.**
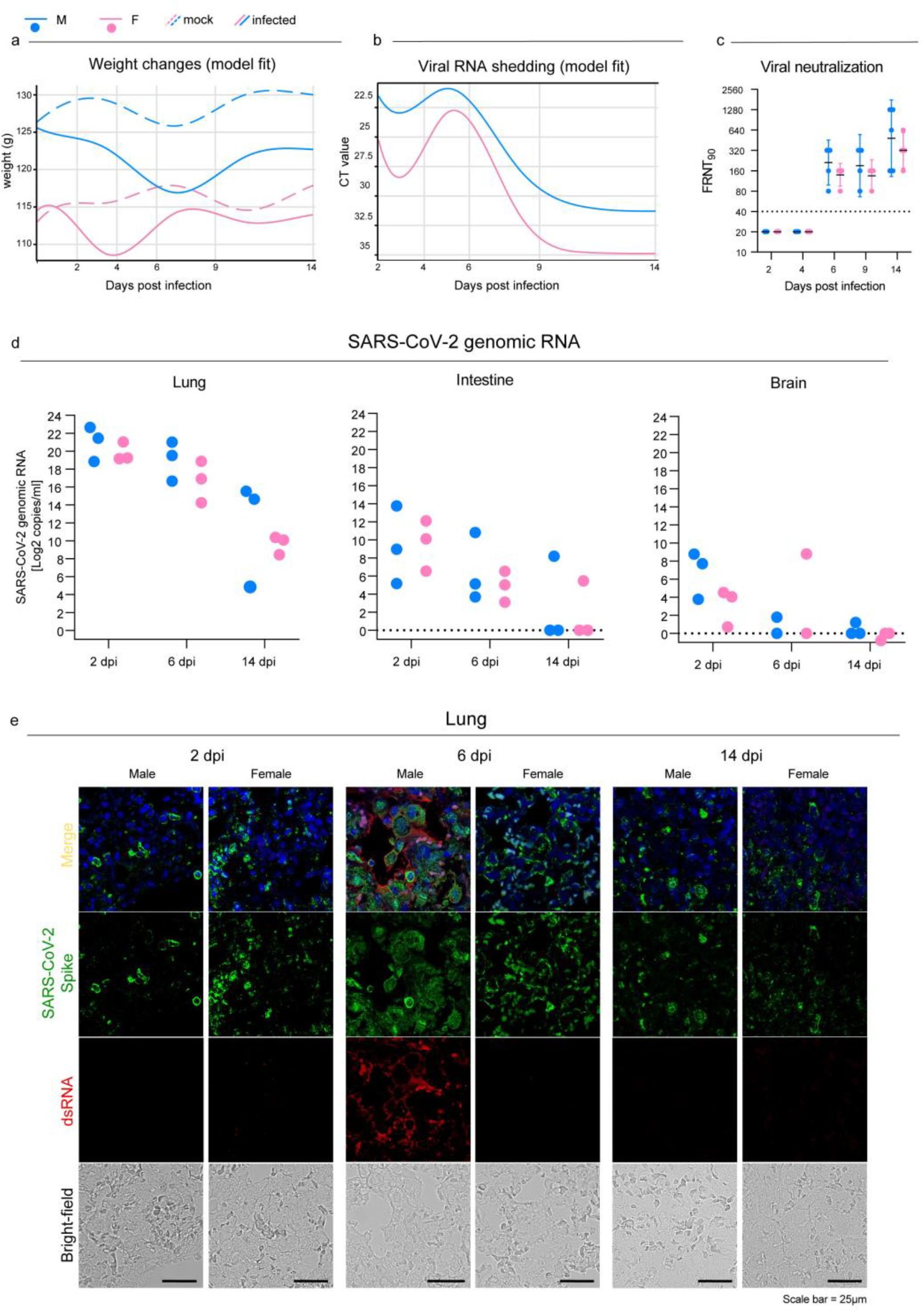
SARS-CoV-2 infection in oropharyngeal swabs, lungs and distal organs. **a**, Statistical model describing body weight changes in mock and infected male (Mock M, M) and female (Mock F, F) Syrian hamsters. **b**, Statistical model describing rRT-PCR results trend of RNA extracted from oropharyngeal swabs at 2, 4, 6, 9 and 14 dpi. Figure 1 a-b: infected and control females 2 dpi n = 30; 4 dpi n = 20; 6 dpi n = 20; 9 dpi n = 10; 14 dpi n = 10 infected and control males 2 dpi n = 30; 4 dpi n = 20; 6 dpi n = 20; 9 dpi n = 10; 14 dpi n = 10. See Table S1 for further details. **c**, Focus reduction neutralization test (FRNT), expressed as the reciprocal of the highest dilution resulting in a reduction of the control focus count > 90% (FRNT90). Geometric means (GMT) with 95% confidence intervals (CI) are represented. Dotted lines indicate the limit of detection (LOD). Wilcoxon-MannWhitney test male vs females; 2dpi *P* > 0.99, 4dpi *P* > 0.99, 6dpi *P* = 0.16, 9dpi *P* = 0.63, 14dpi *P* = 0.59. Infected females 2 dpi n = 5; 4 dpi n = 5; 6 dpi n = 5; 9 dpi n = 4; 14 dpi n = 5 infected males 2 dpi n = 5; 4 dpi n = 5; 6 dpi n = 5; 9 dpi n = 4; 14 dpi n = 5. **d**, SARS-CoV-2 viral load as determined by RTddPCR in the lungs, intestines and brains at 2, 6 and 14 dpi; results are expressed as Log2 copies/ml of genomic RNA for graphical comparison among organs. Infected female and male lungs and intestines: 2, 6 and 14 dpi n = 3 each infected female and male brains: 2 dpi n = 3; 6 dpi n = 2; 14 dpi n = 3 each. No statistical analyses were performed on this dataset due to low sample size. **e**, Representative immunofluorescence staining for SARS-CoV-2 Spike glycoprotein (green) and dsRNA (red) in infected male and female lungs. Scale bar = 25μm. All animals were analyzed, representative images are shown.

**Figure 2.**
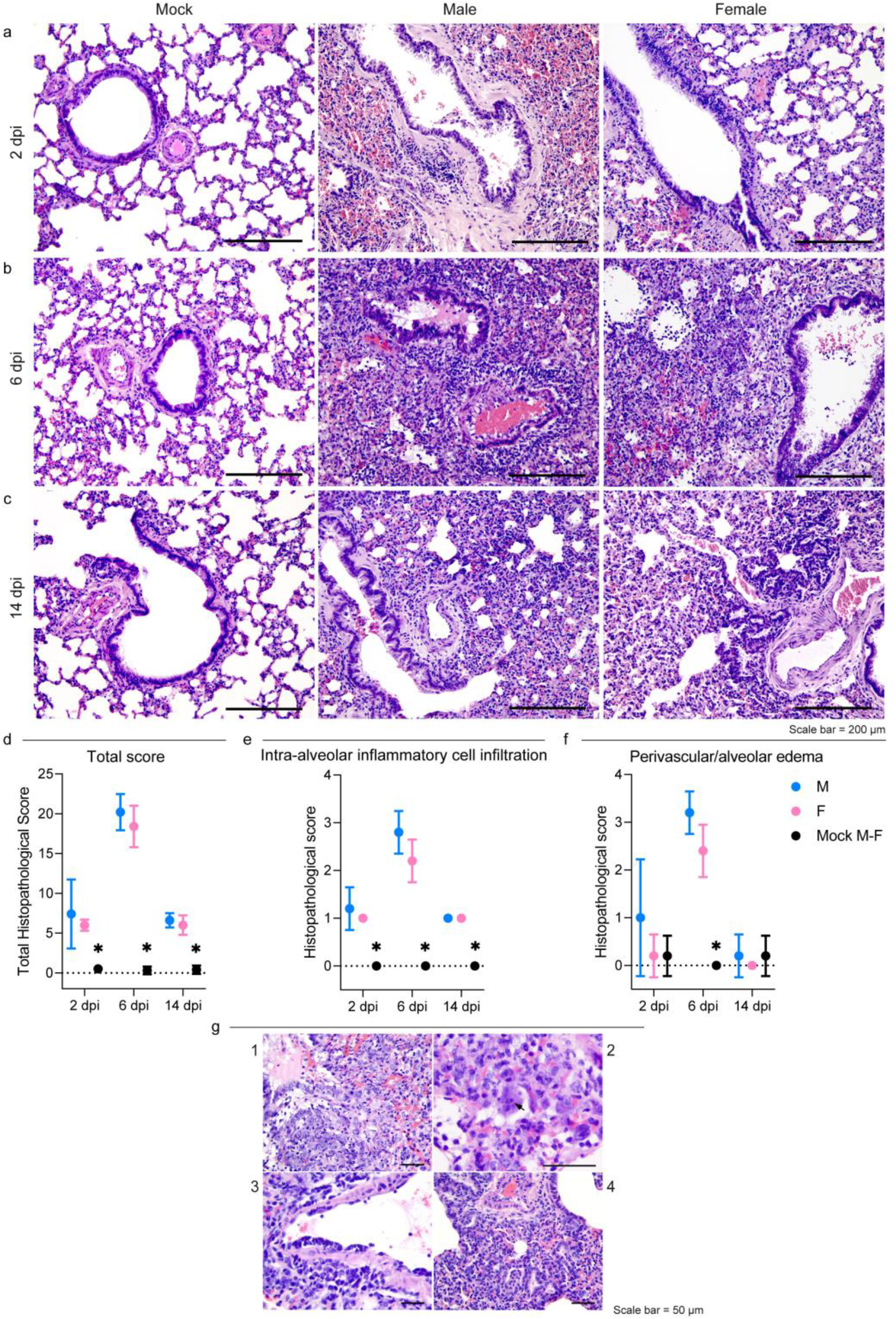
SARS-CoV-2 infection results in severe but rapidly resolving pulmonary lesions in male and female Syrian hamsters. **a, b**, and **c**, Representative images of Syrian hamster lungs collected at 2, 6 and 14 dpi and mock animals. H&E stained sections. Scale bar = 200 μm. All animals were analyzed, representative images are shown. **d**, Cumulative score of lung pathology for nine histopathological assessments in male and female hamsters (Table S2); mean values ± SD are represented. Wilcoxon-Mann-Whitney test males vs females; 2dpi *P* = 0.86, 6dpi *P* = 0.31, 14dpi *P* = 0.72. **e**-**f**, Histopathological scores of intra-alveolar inflammatory cell infiltration (e) and perivascular/alveolar edema (f) in males and female hamsters (Table S2; all animals were analyzed). Mean values ± SD are represented. Wilcoxon-Mann-Whitney test mock animals vs females of total histopathological score (2dpi *P* = 0.0007, 6dpi *P* = 0.0003, 14dpi *P* = 0.0003), intra-alveolar inflammatory cell infiltration (2dpi *P* = 0.0003, 6dpi *P* = 0.0003, 14dpi *P* = 0.0003) and perivascular/alveolar edema (2dpi *P* > 0.99, 6dpi *P* = 0.0003, 14dpi = 0.52). Wilcoxon-Mann-Whitney test mock animals vs males of total histopathological score (2dpi *P* = 0.0007, 6dpi *P* = 0.0003, 14dpi *P* = 0.0003), intra-alveolar inflammatory cell infiltration (2dpi *P* = 0.0003, 6dpi *P* = 0.0003, 14dpi *P* = 0.0003) and perivascular/alveolar edema (2dpi *P* = 0.19, 6dpi *P* = 0.0003, 14dpi > 0.99). Wilcoxon-Mann-Whitney test male vs females of total histological score (2dpi *P* = 0.86, 6dpi *P* = 0.31, 14dpi *P* > 0.72), intra-alveolar inflammatory cell infiltration (2dpi *P* > 0.99, 6dpi *P* = 0.21, 14dpi *P* > 0.99) and perivascular/alveolar edema (2dpi *P* = 0.40, 6dpi *P* = 0.12, 14dpi > 0.99). * indicates a statistically significant comparison. **g**, 1, Severe bronchiolar epithelium and pneumocyte II hyperplasia at 6 dpi; nuclei of proliferating cells were frequently megalic with prominent nucleoli and numerous mitotic figures. 2, A syncytial epithelial cell containing multiple 2-4 µm amphophilic round cytoplasmic viral-like inclusions, in a male hamster at 6 dpi. 3, Lymphomonocytic endothelialitis and perivascular cuffing in a pulmonary venule at 6 dpi. 4, Alveolar bronchiolization with acinar formations and few interstitial lymphoplasmacytic infiltration at 14 dpi. H&E stained sections. Scale bar = 50μm.

**Figure 3.**
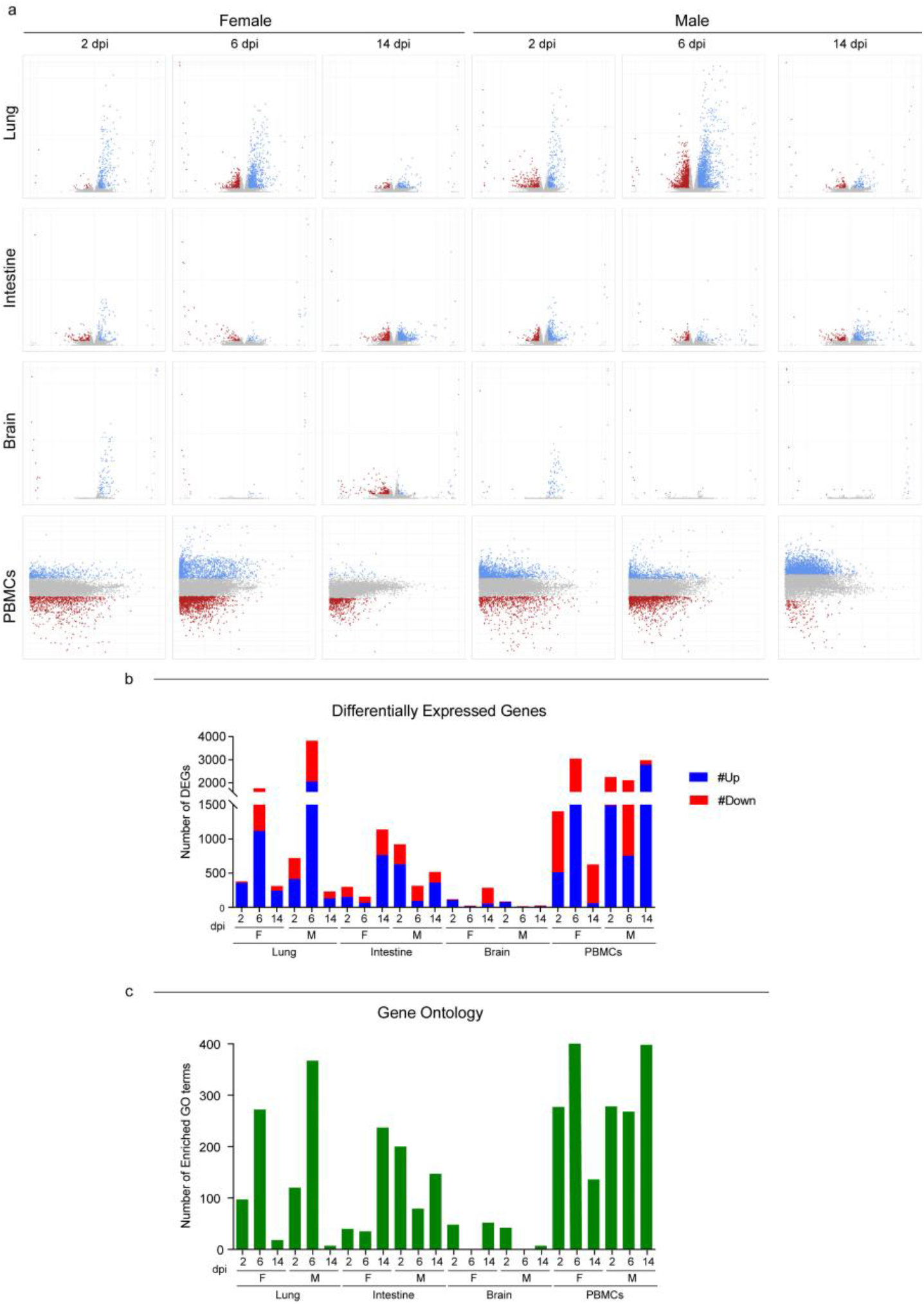
RNA-Seq global expression profiles. **a**, Volcano and MA plot showing differential expression analysis results for lungs/intestine/brains and PBMCs, respectively (blue, up-regulated; red, down-regulated; grey, not significant). A DEG is significant in a comparison when Log2FC ≤ - 1 or Log2FC ≥ 1and FDR < 0.05. For lungs, intestines and brains: x axis = Log2FC; y axis = -LogFDR. For PBMCs: x axis = Log mean expression; y axis = Log2FC. **b**, Number of DEGs for every comparison infected vs mock, done in differential expression analysis; up- and down-regulated genes are shown. See also Table S3-4 for DEGs raw numbers. **c**, Number of enriched GO terms for every comparison infected vs mock done in Gene Ontology enrichment analysis. See also Table S5 for GO terms numbers. Infected female and male lungs and intestines: 2, 6 and 14 dpi n = 3 each; infected female and male brains: 2 dpi n = 3; 6 dpi n = 2; 14 dpi n = 3 each; infected female and male PBMCs: a pool of 5 animals’ blood was analyzed at each time point.

**Figure 4.**
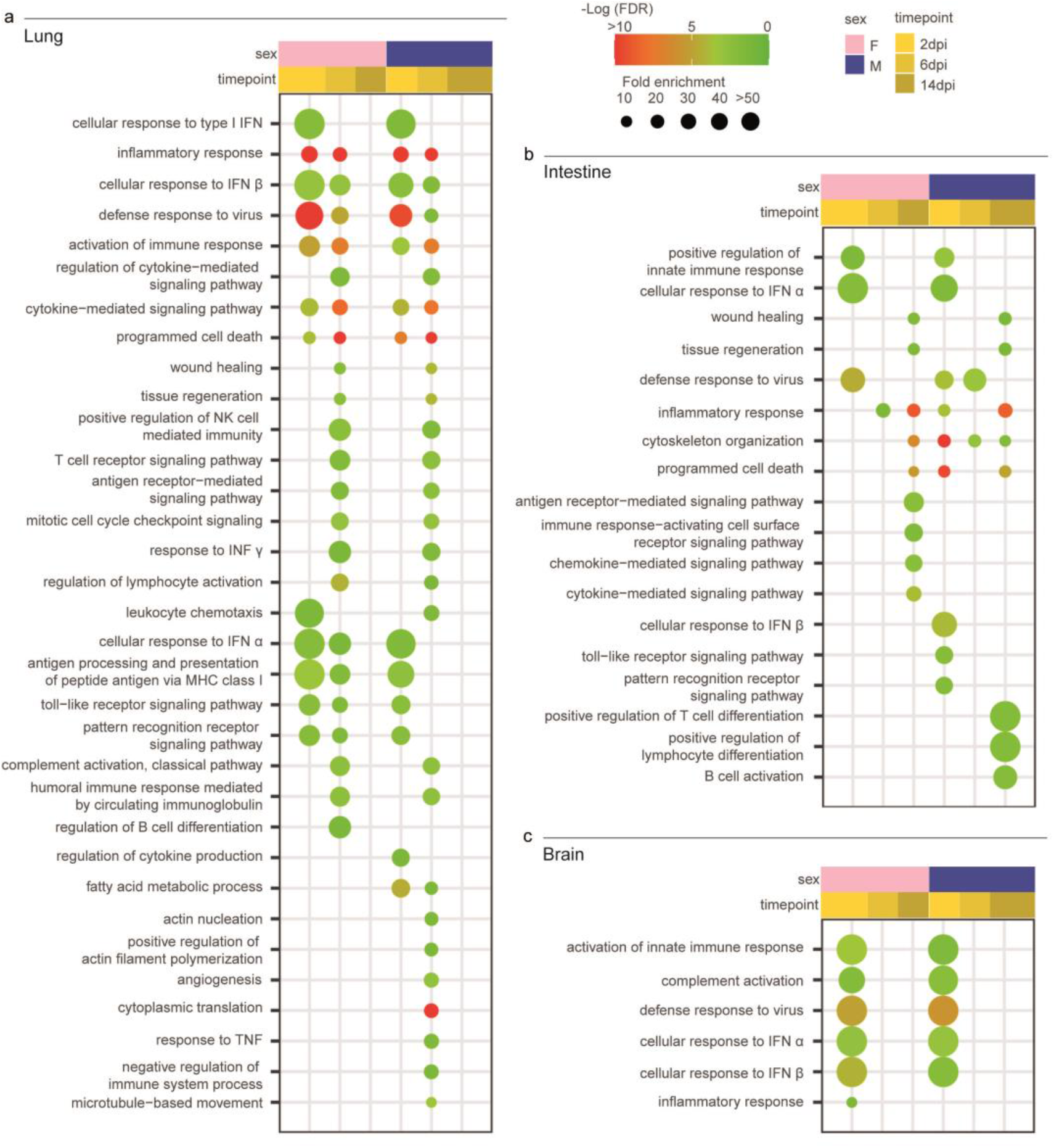
Transcriptomic profile of SARS-CoV-2 infected male and female Syrian hamsters. **a**, Dotplot representing the most specific enriched Gene Ontology (GO) terms related to immunity in lungs. **b**, Dotplot representing the most specific enriched GO terms related to immunity in intestine. **c**, Dotplot representing the most specific enriched GO terms related to immunity in brain. Statistically significant enrichments (FDR < 0.05) are presented and -LogFDR is shown. The absence of the dot means that the indicated GO term is not enriched in that particular sample.

**Figure 5.**
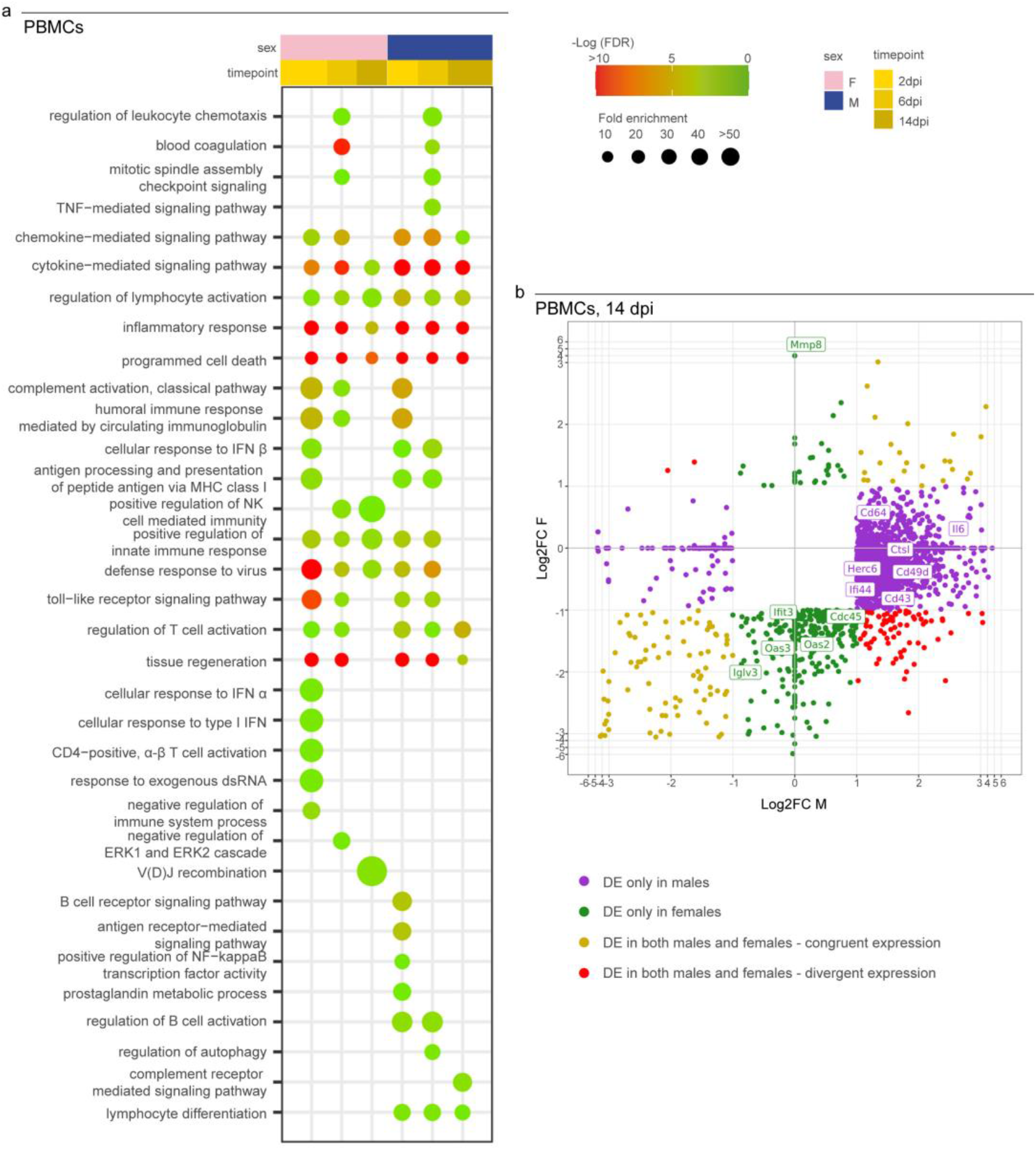
Transcriptomic profile of SARS-CoV-2 infected male and female PBMCs. **a**, Dotplot representing the most specific enriched GO terms related to immunity in PBMCs. Statistically significant enrichments (FDR < 0.05) are presented and –LogFDR is shown. **b**, Scatterplot representing the Log2FC of males and females DEGs in PBMCs at 14 dpi. DE = Differentially expressed. A DEG is significant in a comparison when Log2FC ≤ -1 or Log2FC ≥ 1and FDR < 0.05. The absence of the dot means that the indicated GO term is not enriched in that particular sample.

**Figure 6.**
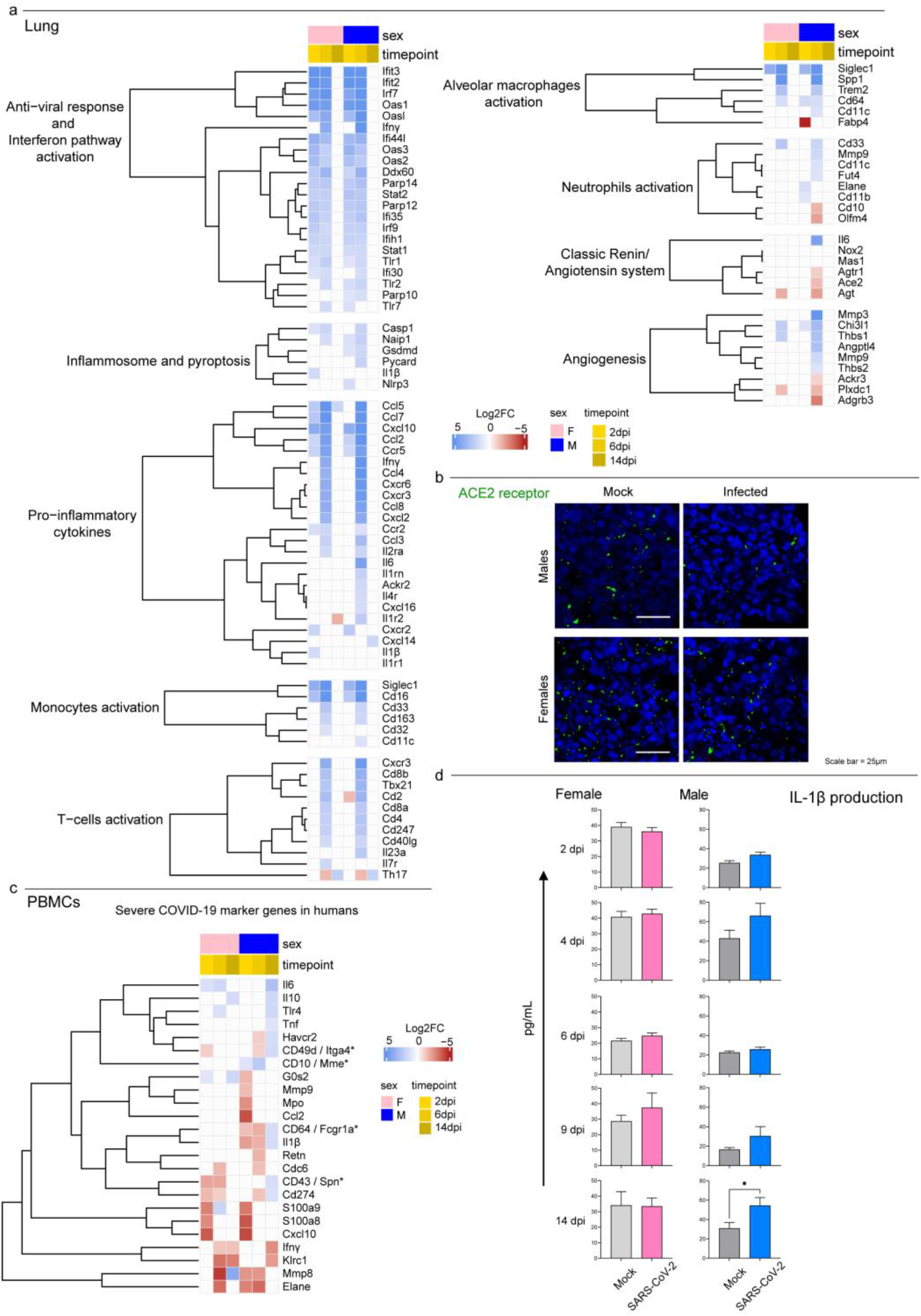
Transcriptomic analysis highlights differences in males and females systemic response to SARS-CoV-2 infection. **a**, Heatmap (Log2FC values of the performed comparisons) of selected genes related to the immune system in the lungs. A DEG is significant in a comparison when Log2FC ≤ -1 or Log2FC ≥ 1and FDR < 0.05. **b**, Immunofluorescence staining for SARS-CoV-2 receptor ACE2 in infected and control males and females lungs at 6 dpi; all animals were analyzed, representative images are shown. Scale bar = 25µm. **c**, Heatmap (Log2FC values of the performed comparisons) of selected genes related to the immune system in the PBMCs transcriptome. *indicates the gene name for hamsters in case it differs from the human ortholog. **d**, Singleplex ELISA levels (pg/mL) for mock and infected male and female hamsters IL-1β. Wilcoxon-Mann-Whitney tests of mock or SARS-CoV-2 M vs F; mean ± SEM is represented (* indicates a statistically significant comparison).

## 3. Results

### 3.1. Infection and seroconversion

Syrian hamsters intranasally infected with the B.1.1.7 SARS-CoV-2 VOC developed no clinical signs, except for a 5% drop in body weight between 2 and 6 dpi, with subsequent recovery (Figure 1a; Table S1 for weight values per sex, status and days post infection). The shedding of viral RNA started at 2 dpi, peaked between 4 and 6 dpi depending on the sex and dropped shortly after. Virus genome was detectable until 14 dpi with high CT values; males showed higher shedding across the whole study period (*P*<0.0001, Figure 1b) with a mean delta of 3.6 CT (Figure 1b; Table S1).

All infected individuals produced detectable neutralizing antibodies against SARS-CoV-2 from 6 dpi, reaching the highest titers 14 dpi (Figure 1c). Geometric mean titers (GMT) were higher in males rather than females, but the difference was not statistically significant.

SARS-CoV-2 established a productive infection in the lungs, with viral RNA detected in all individuals with decreasing viral load over time (Figure 1d). We confirmed these results by showing the presence of the spike protein in the pulmonary parenchyma of all the individual using immunofluorescence. Interestingly, the lungs of male hamsters collected 6 dpi also showed a marked expression of double-strand RNA (dsRNA), which is an indicator of active viral replication, whereas no signal could be observed in the lungs of the females at the same time point (Figure 1e). Mock animals did not stain for any of the tested antibodies, confirming the specificity of reactions. Evidence for SARS-CoV-2 infection in the intestines and brains was far less marked, with low viral load and inconsistent results within the infected groups. On day 14, only one individuals was positive for each group in both organs (Fig. 1d). Coherently with the molecular results, immunofluorescence staining for viral spike glycoprotein and dsRNA was evident only in the intestinal sections of two males at 2 dpi (Figure S1a) and no signal was detected in infected brains at any of the time points analyzed (Figure S1b).

### 3.2. Pathology

Macroscopically, lungs of male hamsters were diffusely consolidated with dark-red coloration on day 6, while multiple dark-red consolidated areas were scattered throughout all the lobes of females. At 14 dpi, we observed few small reddish foci independently of the sex.

Histopathogical changes in the lungs were consistent with a bronchointerstitial pneumonia (Figure 2a-c), with total histopathological score peaking on day 6 in both sexes (Figure 2d; Table S2). At 2 dpi, main histopathological changes consisted in mild-to-moderate alveolar damage, with infiltrates of macrophages and few neutrophils, and vascular hyperemia (Figure 2a). At 6 dpi, extensive and coalescing inflammatory foci with parenchymal consolidation affected more than 75% of the surface in 3 individuals, 50-75% in 5, and 25-50% in 2 females. In all animals, alveolar damage was associated with intense pneumocyte type II and bronchiolar epithelium hyperplasia (Figure 2b and Figure 2g.1). We detected scattered syncytial multinucleated cells in bronchioles and alveolar surfaces that, in one case, contained 2-4 µm amphophilic round cytoplasmic inclusions consistent with viral inclusion bodies (Figure 2g.2). At this stage, edema and infiltration of inflammatory cells (perivascular lymphomonocytic cuffs, alveolar macrophages and neutrophils) were moderate-to-severe, slightly more abundant in males (Figure 2e, f and Table S2). Scattered fibrin exudation in alveolar lumina was detected, with no hyaline membrane formation. Pre- and post-capillary vasculature exhibited plumped reactive endothelium with a sub-endothelial infiltration of lymphocytes, monocytes and rare neutrophils in most animals, consistent with endothelialitis [64] (Figure 2g.3). Infiltrates of inflammatory cells decreased by day 14 when only few lymphocytes, plasma cells and histiocytes surrounding the alveolar ducts were observed (Figure 2c). Alveoli adjacent to terminal bronchioles were multifocally lined by cuboidal cells resembling bronchiolar epithelium (alveolar bronchiolization) [46,65] (Figure 2g.4). There was no evidence of fibroplasia and reparative fibrosis (Figure 2c).

Intestines and brains showed no gross nor histologically detectable lesions (Figure S1c-d).

### 3.3. Host response to SARS-CoV-2 infection

With the purpose of studying male and female host response against SARS-CoV-2, we performed an RNA-Seq analysis on lungs, intestines, brains and peripheral blood mononuclear cells (PBMCs) at three different time points. The comparison of the expression profile of all tissues from infected and mock individuals of the same sex and time point allowed us to quantify and describe the host response in terms of differentially expressed genes (DEGs) (Table S3-S4). In the lungs, the response began at 2 dpi, reached the apex at 6 dpi and was still persistent in the latest time point, with no substantial differences between male and female hamsters. Females PBMCs exhibited the same parabolic curve observed in the lungs, while males elicited a stronger systemic response involving more than 2000 DEGs throughout the study (Figure 3a-b and Table S3-S4). In the intestines and brains, consistently with viral presence and replication, we observed that host response was far less marked. In both organs, females and males followed opposite trends, with DEGs number increasing in females and decreasing in males (Figure 3b). We then employed Gene Ontology (GO) resource to investigate biological processes enriched in SARS-CoV-2-infected Syrian hamsters (Table S5). Except for the intestines of males that showed many enriched GO terms, the number of enriched processes followed the same trend of DEGs in all tissues and sexes (Figure 3c).

To better investigate how Syrian hamster respond to SARS-CoV-2, we focused on GO terms associated to the immune response and correlated biological functions. In the lungs, some GO terms showed the same pattern of enrichment across sexes, being activated in all the infected hamsters at 2 dpi (e.g. “cellular response to type I interferon*”*), 6 dpi (e.g. “T cell receptor signalling pathway” and “response to interferon-gamma”) or both (e.g. “inflammatory response”, “defense response to virus”, “activation of immune response”, “cytokine-mediated signalling pathway”) (Figure 4a). On the other hand, at 6 dpi some GO terms were exclusively enriched in males (e.g. “angiogenesis” and “negative regulation of immune system process”) or exclusively in females (“regulation of B cell differentiation”) (Figure 4a).

In both the intestines and brains, we were unable to find a clear inflammatory pattern in response to the infection with SARS-CoV-2 (Figure 4b-c).

In the intestines we found few GO terms related to the immune system. At 2 dpi, “positive regulation of innate immune response”, “defense response to virus” and “cellular response to interferon-alpha”, were enriched in both sexes, while “cellular response to interferon-beta” and “toll-like receptor signalling pathway” were specifically enhanced in males. Only three GO terms of very general means (e.g. “inflammatory response”) were enriched at 6 dpi, while at 14 dpi we detected GO terms related to lesions recovery, such as “wound healing”, and “tissue regeneration”. At this time point, we noted few sex-specific enriched terms, such as “positive regulation of T cell differentiation”, “lymphocyte differentiation” and “B cell activation” in males and “antigen receptor-mediated signalling pathway”, “chemokine-mediated and cytokine-mediated signalling pathways” in females (Figure 4b).

In the brains, few GO terms were enriched in both sexes exclusively at 2 dpi, including “activation of immune response”, “complement activation”, “defense response to virus”, “cellular response to interferon-alpha and -beta” (Figure 4c).

As a major novelty of this study, we analyzed the immunological profile of PBMCs to investigate the Syrian hamster systemic activation of the immune system, searching for potential similarities with human severe COVID-19 cases. We observed the activation of the immune response in both sexes in all the three time points, as expressed by longitudinal enrichment of related GO terms such as “cytokine/chemokine-mediated signalling pathway”, “regulation of lymphocyte activation”, “inflammatory response”, “programmed cell death” and “defense response to virus” (Figure 5a). Other GO terms enriched in both sexes during the experiment at any time point, including “complement activation”, “antigen processing and presentation of peptide antigen via MHC class I”, “positive regulation of innate immune response” and “toll-like/pattern recognition receptor signalling pathway”. Some GO terms were enriched in a sex-specific manner, such as “regulation of autophagy” and “lymphocyte differentiation in males”, or “cellular response to interferon-alpha” and, *“*alpha-beta T cell activation” in females. In particular, we observed major differences between females and males at 14 dpi, when 98% of the upregulated genes (2676/2723) were male-specific and 60% of down-regulated genes (401/672) were female-specific (Figure 5b and Table S4).

### 3.4. Syrian Hamster as immunological model for COVID-19

To investigate whether hamsters display the typical immunological profiles described in COVID-19 lungs, we evaluated the expression levels of 100 genes associated to a severe human condition in previous studies [13,65–71] (Figure 6a). Syrian hamsters activated an Interferon-I (IFN-I)-mediated cell-specific response to the virus at 2 dpi, as shown by the up-regulation of many interferon stimulated genes (ISGs). This included genes coding for IFIT proteins (e.g. *Ifit2, Ifit3*), members of the OAS family (e.g. *Oas1, Oas2, Oas3, Oasl*), interferon regulatory factors (e.g. *Irf7* and *Irf9*) and several genes involved with cellular mechanisms of antiviral response (e.g. *Ddx60, Parp12* and *Parp14*). Specific immune response increased in both sexes at 6 dpi, with upregulation of 58 and 50 out of 100 target genes for males and females, respectively. SARS-CoV-2-infected animals promoted immune cell recruitment with complement activation, immunoglobulin-mediated response, and strong upregulation of pro-inflammatory cytokines (e.g. *Ccl2, Ccl3, Cxcl10, Il6* and *Ifnγ*) moreover, we observed the activation of the genes involved in monocyte (*Cd33, Cd16* and *Siglec1*) and T-cell activation (*Tbx21, Cd40lg, Cd4, Cd8a* and *Cd8b*). At 6 dpi males also up-regulated genes associated with active neutrophils recruitment (e.g. *Mmp9, Cd11c, Fut4* and *Elane*) and angiogenesis (e.g. *Mmp3, Thbs1* and *Angptl4*) (Figure 6a). Interestingly, male hamsters downregulated both SARS-CoV-2 receptor *Ace2* and its receptor *Agtr1* genes. Of note, immunofluorescence staining for ACE2 expression confirms the sex-specific reduction of the receptor in lung tissue compared to mock controls (Figure 6b). Hamsters of both sexes shut down almost completely the specific pulmonary immune response by day 14, with no individual perpetuating the immune exasperation and inflammation typical of severe COVID-19.

Among 32 key immunological genes associated to a severe COVID-19 systemic pathology in humans, 24 were differentially expressed in hamsters’ PBMCs in at least one case (Figure 6c). Our results present a male-biased up-regulation of genes associated with immature neutrophils activation (e.g. *Cd49d, Cd274, Tlr4* and *Cd43*) and pro-inflammatory cytokines associated with the cytokine storm (*Il1β, Il6* and *Tnf*). In this context, the longitudinal monitoring of pro-inflammatory IL-1β and IL-6 revealed their low release in the serum in response to SARS-CoV-2 infection, with exclusive increase in circulating levels of IL-1β in male hamsters at 14 dpi (Figure 6d; Figure S2).

## 4. Discussion

Following the declaration of the COVID-19 pandemic by the WHO in March 2020, both the scientific community and health authorities were on the frontline for the development of control measures to limit the spread of the infection and mitigate disease severity. To achieve this goal translational animal models were used to elucidate the pathogenesis of the disease and to rapidly assess the efficacy of prophylactic and therapeutic agents. However, in order for scientists to select the best animal models for their studies, it is crucial to characterize in which way a species can mimic the host-pathogen relationship between humans and SARS-CoV-2. In this study, we provide a comprehensive description for the Syrian hamsters that, long before the emergence of SARS-CoV-2, has been extensively used as an animal model to study other zoonotic emerging diseases, including bunyaviruses, arenaviruses, henipaviruses, flaviviruses, alphaviruses, filoviruses, as well as the coronaviruses SARS-CoV and MERS-CoV [72]. We performed experimental infections using SARS-CoV2 B.1.1.7 strain or Alpha VOC, isolated in Italy. In our study, we successfully infected all the hamsters, detecting RNA of SARS-CoV-2 in tissues and oropharyngeal swabs from day 2 to 14 and specific antibody response by day 6, supporting earlier evidences [38,40,42]. In addition, we confirmed the presence of the antigen within the pulmonary tissue up to 14 dpi through immunofluorescence, using a specific antibody directed towards the spike protein. Unfortunately, our choice to store samples only in RNA later and formalin driven by the need of securing the quality of molecular and histological analyses, hampered any isolation attempt to test the viability of the virus within different tissues. However, based on other studies, no viable virus is present in the lungs after day 7 [42] and this makes us assume that our findings are associated to the presence of residual RNA/proteins rather than to active infection. Indeed, we implemented a specific immunofluorescence technique to detect dsRNA, a replicative intermediate of many RNA viruses, coronaviruses included [48,50]. Interestingly, we found positive staining for dsRNA only in male lungs at 6 dpi, suggesting a sex-related difference in the replication rate. However, such a difference did not translate into an evident increase in the total virus using ddPCR. In this sense, paired immunofluorescence and molecular investigations performed at intermediate timepoints might have helped to elucidate the dynamics of viral infection and replication within the pulmonary tissue of female and male hamsters.

SARS-CoV-2-infected hamsters developed a moderate-to-severe bronchointerstitial pneumonia mimicking histological patterns observed in COVID-19 patients (i.e. diffuse alveolar damage –DAD-, interstitial and intra-alveolar influx of macrophages/neutrophils and pulmonary vascular endothelialitis), as previously described [38,40,43,44,73]. Compared to the pneumonia described in human COVID-19, DAD was milder and unevenly distributed; in addition, we did not notice the formation of hyaline membranes. Albeit some studies confirmed DAD in hamsters using the original USA-WA1/2020 isolate [39], our findings are in line with the current literature that supports the evidence of a milder disease in this animal model, especially considering that most human findings derive from patients died of severe COVID-19 [38,42,74–76].

Despite lung damage, hamsters showed no clinical signs but a significant loss in body weight that resolved spontaneously by day 14 post-infection. This result is consistent with previous reports [31], although few studies also described symptoms such as lethargy, ruffled fur, hunched back posture and rapid breathing [38,39], a difference that might be related with the virus (i.e. titer and route of inoculum or viral strain) [40,77], the hamsters (i.e. age) [40,78] or a combination of both [40]. While it is known that prey species such as hamsters mask their sickness when they perceive a threat, such as the presence of humans [79], these data suggest that the disease in hamsters mostly resembles that found in humans with mild COVID-19 symptoms. In humans, severe COVID-19 is associated with tissue damage due to an exacerbated inflammatory response [80] and multi-organ failure as secondary effect of systemic activation and exhaustion of the immune system [66,81], or due to viral spread outside the respiratory system [82,83]. In our study, we investigated the viral spread and the hamsters’ immune response at local and systemic level, in order to evaluate the differences between our animal model and severe cases of COVID-19 in humans.

Hamsters mostly responded to SARS-CoV-2 in the lungs within the first week, with subsequent silencing by the end of the experiment that paired the recovery from the clinical disease, the clearance of the infection and the repair of pathological lesions. Most DEGs and GOs were associated to the immune response and related biological functions, including the activation of IFN-I alpha and beta, as previously described [42,44]. These molecules are crucial for effective antiviral response because they counteract viral replication in infected cells and cell-to-cell spread, enhance antigen presentation, and promote the development of the adaptive immune response [84,85]. Despite the induction of interferons being dampened after the infection with SARS-CoV-2 compared to other viruses such as Influenza A [14,86], IFN-I signalling influences the severity of COVID-19 in humans. Alterations in TLR3-dependent and TLR7-dependent type I interferon induction, the presence of autoantibodies to interferon and, in general, the reduced induction of local and systemic interferon responses against SARS-CoV-2 infection lead to severe manifestations [14]. Indeed, restricted IFN-I response might promote longstanding active viral replication, excessive production of pro-inflammatory cytokines and influx of neutrophils and monocytes, which act as further sources for pro-inflammatory mediators and promote greater tissue damage [80]. In this context, it is likely that the early and powerful induction of IFN-I related genes that we described in hamsters promotes fast viral clearance in the lungs and tissue structure restoration, preventing severe manifestations of the disease and the systemic spread of the virus in this animal model. Our data show that, similarly to humans, also hamsters respond to the infection with local inflammation, recruitment of immune cells, activation of the complement and immunoglobulin-mediated response, and release of pro-inflammatory cytokines. However, such a response is contained in this animal model and shut down by day 14 post-infection. Furthermore, PBMCs RNA-Seq data showed a modest systemic response in hamsters that resolves within two weeks, with the activation of the interferon pathway, innate cell recruitment and activation of lymphocytes B and T and immunoglobulin−mediated immune response. A modest increase of the circulating levels of pro-inflammatory cytokines further corroborates previous studies [87,88] and highlights another crucial difference with patients suffering from complicated COVID-19 that present almost 3-fold higher levels of pro-inflammatory cytokine IL-6 compared to patients with an uncomplicated form of the disease [89]. Overall, our data suggest that hamsters do not suffer of any dysregulation of the immune system that might determine severe COVID-19 in humans.

Consistently with the low systemic activation of the immune system that in humans promotes tissue damage in peripheral districts, we discovered that there were no histopathological lesions in the intestines and brains of the hamsters. In addition, our ddRT-PCR data support other studies in showing the limited spread of SARS-CoV-2 outside the respiratory tract in this species [40]. This data is in line with the current literature, including minimal spread to the kidneys and heart [42]. The lower or absent systemic infection in hamsters compared to humans, where the virus can spread to the digestive tract, the brain, the heart, the kidneys, the sweat glands of the skin and the testicles [82,83] further explains the fewer complications seen in this model. Interestingly, we found positive staining in immunofluorescence for the spike and dsRNA, thus supporting replication of the virus in the intestines, with transcriptomic analyses showing weak and generic immune response. While the lack of studies on the transcriptome of human intestines during COVID-19 prevents us from making significant comparisons with our animal model, infection of human small intestinal organoids resulted in much higher transcriptomic signal [90]. On the other hand, the minimal alterations shown in our analyses could simply result from enterocytes sloughing following fasting and weight loss.

In our study, all data supported the assumption that infection with SARS-CoV-2 has more severe consequences in male hamsters. Indeed, males developed more diffuse and severe lung lesions, characterized by higher scores of infiltration of inflammatory cells and edema, which may have resulted in the more obvious pathological manifestations. Thanks to the combination of several approaches, our study allowed us to investigate the possible causes and consequences of such a difference. Of note, we found that in the lungs, males display a higher differential expression of genes associated both with activated neutrophils and alveolar macrophages and with the release of pro-inflammatory cytokines associated to ARDS, such as *Il-6, Cxcl10* and *Ifnɣ*. This sex-based difference has been evidenced in human COVID-19 cases [91–94] but it had not been previously reported for animal models, where the transcriptome of infected hamsters was mainly investigated using RT-qPCR rather than RNA-Seq analysis [78,95,96]. Another peculiarity of male hamsters standing out from our data is the differential expression of genes promoting angiogenesis (e.g. *Mmp3, Thbs1* and *Angptl4*), that might explain sex-driven differences in the pulmonary lesions. Finally, male hamsters downregulate both *Ace2* and its receptor *Agtr1* on day 6, a feature that we were able to identify using transcriptomic analyses and to confirm through immunofluorescence, showing a decreased level of the receptor within the pulmonary tissue between non-infected and infected animals. As the receptor gets endocytosed together with the virus during cellular infection, this difference might be due to a higher level of infection and replication of SARS-CoV-2 or to an increased cell death and apoptosis in males. In this context, we found a general enrichment of the programmed cell death process in lungs between 2 and 6 dpi, but no differences between females and males (Figure 4a). On the other hand, we observed a high viral load by ddRT-PCR and a peculiar staining for dsRNA in the lungs of male hamsters, corroborating a possible link between the regulation of *Ace2* and the replication of the virus. Other than being SARS-CoV-2 cellular receptor, ACE2 has the physiological function of inactivating angiotensin II (AII) molecules produced by ACE, known for its vasoconstrictive activities and, crucially, for acting as a potent pro-inflammatory cytokine [80]. As a further notice, increased level of AII can also exacerbate IL-6 signalling. In this context, several cytokine storm cytokines [81,97] as *Il-6, Il-1β* and *Tnf* and genes associated with immature neutrophils activation (e.g. *Itga4, Cd274* and *Spn*) were specifically up-regulated in PBMCs from male hamsters only. Similarly, males showed a peculiar increase of serum levels of IL-1β at 14 dpi that was not observed in females, thus suggesting a possible re-acerbation of the systemic inflammation.

These evidences further corroborate a sex-mediated difference in the pathology of COVID-19 in hamsters that could provide useful insights to understand similar evidences in humans. Indeed, studies worldwide support that more men than women require intensive care or succumb to the disease [16]. While it has been suggested that social and behavioural differences between genders might influence the progression of COVID-19, our data support the role of the biological sex. Finally, the longitudinal assessment of oropharyngeal swabs demonstrated that, while showing akin kinetics, males eliminate more virus, suggesting sex-driven differences also in the epidemiology of the pandemic.

## Conclusions

Our study provides a comprehensive evaluation of the Syrian hamster as animal model for COVID-19. Overall, we confirmed that the infection with SARS-CoV-2 shows similar pathways both in humans and hamsters, which proved to be an excellent model to test the efficacy of prophylactic biologicals, such as vaccines, and to quickly assess the phenotypic changes of new VOCs [31]. As a matter of fact, our clinical and histopathological data suggest a decreased pathogenicity of the Alpha VOC compared to the original strain, which have been found to induce DAD and cause more severe symptoms in the hamster [38,39]. More recently, the experimental infection of hamsters was also effective in showing the decreased pathogenicity of the VOC Omicron upon emergence, which confirmed preliminary epidemiological data obtained from the human population [43]. In addition, Syrian hamsters have been successfully used to investigate the transmission of SARS-CoV-2 between cohoused contact hamsters [98], a feature that often requires the use of ferrets, a model that is associated with higher costs and logistical challenges [31]. While the fast recovery of the animals and clearance of the virus observed in our study would suggest otherwise, the Syrian hamster was also found as an effective model to evaluate the long-term effects of SARS-CoV-2 infection, collectively known as “long-COVID syndrome” or post-acute sequalae of COVID-19 (PASC) [99,100], which is still a very heterogeneous and poorly understood condition in humans. Indeed, Frere et al., [42] showed that hamsters infected with the original strain WA1/2020 display transcriptional alterations in the striatum brain and persistent inflammatory profile in the olfactory bulb more than one month after infection, which could be associated with cognitive impairments, depressive disorders and behavioural changes seen in patients suffering from long COVID [42].

On the other hand, our study underlines that hamsters only mimic mild-to-moderate COVID-19 and neither replicate the exacerbation of the immune response nor the systemic spread. So far, the hamster model failed to recapitulate the most severe human symptoms regardless of the variant used in the experiments, suggesting that our data could be at least partially extrapolated outside the specific case of the VOC Alpha [31]. In this context, hamsters should be used with caution for more detailed studies on COVID-19 and to evaluate therapeutic agents dampening the immune response. Among other species, African green monkeys are currently the best model to mirror severe disease manifestations seen in humans, notably ARDS [31]. Most recently, mice model engineered or engrafted with human tissue also proved effective in mimicking physiological features of human infection, allowing for detailed pathological studies [31]. However, most of these models have several constraint for widespread use, such as lower availability, higher costs, and difficulties in handling and managements within experimental settings. It is also crucial to mention the increasing role of alternative models, among which organoids obtained by human stem cells have emerged as powerful tools able to bridge the gap between cell lines and animal models and to scale-up to high-throughput screens [101]. These systems have been widely used to study SARS-CoV-2 infection, including viral tropism, host response and drug discovery of different VOCs [101]. Future perspectives include the production of organoids with diverse genetic backgrounds to explore the impact of host genetics on disease progression, response to vaccines and therapeutic agents [102].

As a final note, we were able to observe a significant difference between female and male hamsters that should be taken into account when designing any experimental study. While this feature is possibly not peculiar to SARS-CoV-2 infection, the sex-biases of animal experiments has long represented a critical aspect of translational medicine[103]. Fortunately, researchers, funders and policy makers unanimously acknowledge the need for a change; research projects that include both sexes and analyses of data by gender – as in the present study - are becoming more and more popular. In turn, we believe that animal models will progressively become important not only to describe the pathological pathways of a disease but also to grasp differences related to biological sex.

## Ethical statement

Animal studies were performed in compliance with directive 2010/63/EU of the European Parliament and of the Council of 22 September 2010 on the protection of animals used for scientific purposes. The experimental design was approved by IZSVe ethical board and by the Italian Ministry of Health, under permit n. 1167/2020-PR. In accordance with the 3Rs principle (Replacement, Reduction and Refinement), we used the minimum number of animals that secured statistically sound results and provided best housing and environmental enrichment. Briefly, individual housing exceeded the minimum surface required and agreed with the ecology, behaviour and biology of the species. Temperature, humidity and light-dark cycles were fixed (21 ± 3 °C, 50 ± 10%, lights off: 07:00 AM–07:00 PM) and monitored throughout the study. All animals had *ad libitum* access to food and water throughout the entire study. Environmental enrichment consisted of gnawing blocks, nesting material and extra sunflower seeds three times a week. We guaranteed daily monitoring of animals’ health and comfort and established a humanitarian threshold to avoid unnecessary suffering. Animals were bred *in house* at the Istituto Zooprofilattico Sperimentale delle Venezie, under permission N n°2020/0095 granted by the municipality of Legnaro on August 2020.

## Acknowledgments

The authors would like to thank Francesco Bonfante for having kindly provided the viral strain used in this study. We acknowledge Massimo Boldrin, Franco Mutinelli, Maria Augusta Bozza and the whole team working at the IZSVE animal facilities for their support in breeding and managing the hamsters. We also thank Annalisa Salviato, Alessia Schivo, Miriam Abbadi, Lorena Biasini, Sara Petrin and Arianna Peruzzo for their training, support and precious suggestions on wet lab techniques for transcriptomic analyses. We also thank Giorgia Monetti for processing fixed organs and providing tissue slices for immunofluorescence and histological analyses. Finally, we are grateful to Francesca Ellero for her English edits on the manuscript.

## Author Contributions

Funding acquisition and project administration: C.T., P.D.B and S.L.; supervision: S.C. and P.D.B.; resources: A.O., M.V., I.M., C.T. and P.D.B.; study design: M.C., G.Z., M.M., P.D.B., and S.L.; animal experiments: M.C., M.Z., P.D., P.D.B. and S.L.; histology: Gr.F. and M.V.; molecular analyses: M.C., P.D., I.B. and V.P.; immunofluorescence: M.C.; transcriptomic analyses: M.C. and G.Z.; serology: A.B. and M.P.; ELISA: Gi.F.; statistics: M.M.; data interpretation: M.C., G.Z., Gr. F., Gi.F., S.C., P.D.B. and S.L.; writing (first draft): M.C., G.Z., Gr.F, Gi.F., M.M., V.P. and S.L.; writing (revision): all authors; Figures: M.C., G.Z., Gr.F., Gi.F., M.M. and S.L.

