## Supplementary material for "Host response of Syrian hamster to SARS-CoV-2 infection, including differences with humans and between sexes"

**Supplementary methods**

**Sample collection and storage.** Scheduled necropsies of Syrian hamsters (*Mesocricetus auratus*) were performed at 2, 6 and 14 days post infection (dpi), involving 5 individuals per group at each time point. We performed terminal blood collection and collected the lungs, brains and intestines, preserving half of each sample in 10% neutral-buffered formalin and the other half in RNA later (Thermo Fisher®), respectively used for histological examination and molecular analyses. In order to maximize the homogeneity between samples, we paid particular attention in collecting two equal parts and devolving the same portion of each organ to perform each analysis (i.e. left or right lung or brain hemisphere and proximal versus distal portion of the intestine). For histological evaluations, the intestines were opened longitudinally, washed in sterile PBS and wrapped. Formalin-fixed lungs, brains and intestines were routinely processed and paraffin-embedded for histological examination. Samples and oral swabs were collected in RNA later and stored for 24-48 hours at 4°C, then the medium was removed and organs/swabs were frozen dried at -80 °C until further analyses.

**PBMCs isolation.** Immediately after euthanasia, we performed a terminal blood collection through intracardial sampling using S-Monovette® K3 EDTA syringes (Sarstedt) for the isolation of peripheral blood mononuclear cells (PBMCs). In order to obtain a sufficient volume for effective isolation, we mixed in equal volumes the blood from five individuals at each time point for each experimental group; one volume of sterile PBS was added to the blood samples. Pooled blood samples were layered onto 15 ml of Ficoll-Paque Plus (GE healthcare) and centrifuged at 400 g for 60 min at room temperature, after which the plasma layer was discarded. The fraction containing PBMCs was collected into a new 50 ml Falcon and washed with 3 volumes of PBS by centrifugation at 300 g for 8 min. Pellet was suspended in PBS and centrifuged again at 300 g for 8 min. PBMCs were then collected in complete medium (RPMI-1640, 1% Pen-Strep, 10% FCS; Gibco, ThermoFisher Scientific), and then both cell number and viability were assessed using Trypan Blue (ThermoFisher Scientific). Cell freezing medium (40% complete medium, 50% FCS and 10% DMSO) was slowly added to PBMCs on ice and the mixture was then cryopreserved in liquid nitrogen.

**Immunofluorescence on tissue sections.** Tissue sections 4 µm-thick were re-hydrated and incubated in citrate buffer 0.01 M at pH 6 and 95°C for 20 minutes to retrieve antigens. We then permeabilized the tissues using PBS additioned with 1% Triton X-100 for 20 min at room temperature, saturated aspecific sites using Blocking Buffer (BSA 5% in PBS 0.1% Triton [PBSt]) for 1 hour, and incubated the slides overnight at 4 °C with the primary antibody. The following day, samples were incubated for 2 hours in the dark with the specific secondary antibody conjugated with a fluorophore that was chosen depending on the primary one and previously diluted in 1% BSA in PBSt. Fluoroshield with DAPI (Sigma) was used as mounting medium in order to label the nuclei of the cells, thus allowing a better visualization, interpretation and counting. Images were acquired with Leica TCS SP8 confocal microscope equipped with a CCD camera at 63X enlargement using LAS AF 2.7.3.9723 software and analyzed using ImageJ.

**Table S6.** Primary and secondary antibodies used for immunofluorescence.

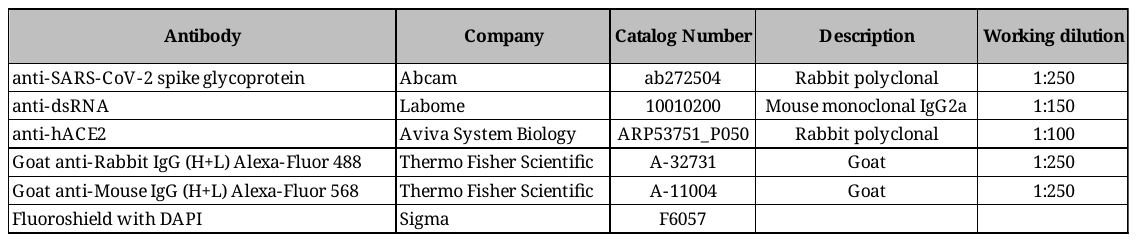

**RNA extraction.** To determine the salivary shedding of virus over the infection period, we rehydrated oropharyngeal swabs with 500 µl PBS and purified nucleic acids from 200 µl suspension using the MagMAX™ Pathogen RNA/DNA Kit (Applied Biosystems™) on a KingFisher™ Flex Purification System (ThermoFisher Scientific; Low-cell-content protocol). To quantify SARS-CoV-2 in target organs, approximately 25 mg of lung, brain and intestine tissues collected from three infected males and three infected females per time point were homogenized in RLT Lysis buffer (Qiagen) using the TissueLyser (Qiagen). Total RNA was isolated using the Rneasy Mini kit (Qiagen) and contaminant DNA removed with the Rnase-Free Dnase Set (Qiagen).

**Qualitative rRT-PCR and RT-ddPCR.** We investigated the presence of viral RNA in oropharyngeal swabs of all the infected and control animals by rRT-PCR using 5 µl RNA, 0.4 µM each primer, 0.2 µM probe and nuclease-free water, up to 22 µl. The thermal cycling conditions were as follows: 50˚C for 30 min, 95˚C for 10 min, 45 cycles of 95˚C for 15 sec and 60˚C for 1 min. The quality of the sample was verified by rRT-PCR targeting β-actin mRNA using 2 µl RNA, 0.4 µM of each primer, 0.2 µM of probe and nuclease-free water, up to 25 µl. The thermal cycling conditions were as follows: 50˚C for 30 min, 95˚C for 10 min, 45 cycles at 95˚C for 15 sec and 56˚C for 30 sec. Both rRT-PCR reactions were carried out using the AgPath-ID one-step RT-PCR Reagents (Applied Biosystems) on a CFX96 Deep Well real-time PCR system (Bio-Rad) in separate tubes. We estimated the number of SARS-CoV-2 genomic RNA copies in the lung, intestine and brain tissues of the individuals randomly selected for the RNA-Seq analysis by RT-ddPCR using the One-Step RT-ddPCR Advanced Kit for Probes (Bio-Rad). Briefly, 5 µl normalized RNA were added to the reaction mixture containing 5 µl supermix, 1 µl 300 mM DTT, 0.9 µM of each primer, 0.25 µM of probe and nuclease-free water, up to 22 µl. To verify the quality of extracted RNA, the β-actin mRNA was co-amplified in the reaction mix. Droplets were produced with a QX200 Droplet Generator (Bio-Rad), according to the manufacturer’s instructions. Amplification was carried out on a C1000 Touch Thermal Cycler (Bio-Rad), under the following cycling conditions: 50˚C for 60 min, 95˚C for 10 min, 50 cycles of 95˚C for 30 sec (ramp rate 2°C/sec) and 60˚C for 1 min (ramp rate 2°C/sec), 98˚C for 10 min. Droplets were scanned with the QX200 Droplet Reader (Bio-Rad) and data analyzed with the QX Manager Software Standard Edition, Version 1.2 (Bio-Rad). Samples with at least 10,000 events were taken into consideration. To increase assay sensitivity in intestine and brain specimens, samples were run in duplicate or triplicate in order to account for a higher number of events.

**RNA-Seq analyses.** Three randomly chosen individuals per experimental group were used for almost all comparisons. For PBMCs, we analyzed a single pooled sample for each group, representative of all five individuals; for brains at 6 dpi only 2 female and male samples were analyzed as, after extraction, they uniquely reached RNA minimum concentration and quality for RNA-seq analyses. The final sample set included 116 specimens. Analyses were preceded by a quality check of RNA in the samples using Agilent RNA 6000 Nano kit, considering as acceptable RIN values ≥ 8. Libraries were prepared from 250-700 ng of total RNA. We filtered raw data by clipping the library adaptors and trimming low-quality ends with trimmomatic v0.39 [1] and we removed remaining reads shorter than 50 bp. To define differentially expressed genes (DEGs) between infected and mock males and females at each time point, we applied DeSeq2 package v1.20.0 [2], using FDR < 0.05 and |Log2FC| ≥ 1in lungs, intestines and brains. For PBMCs, which consisted of a single pooled replicate for each condition, we identified DEGs with GFOLD v1.1.4 [3] using |GFOLD value| ≥1 as reliable Log2 fold change value. GO enrichment was computed by Fisher’s exact test and p-values were adjusted through the Benjamini–Hochberg correction, considering as significant an FDR <0.05. Enrichment scores were computed as -Log(FDR).

**Supplementary figures and figure legends**

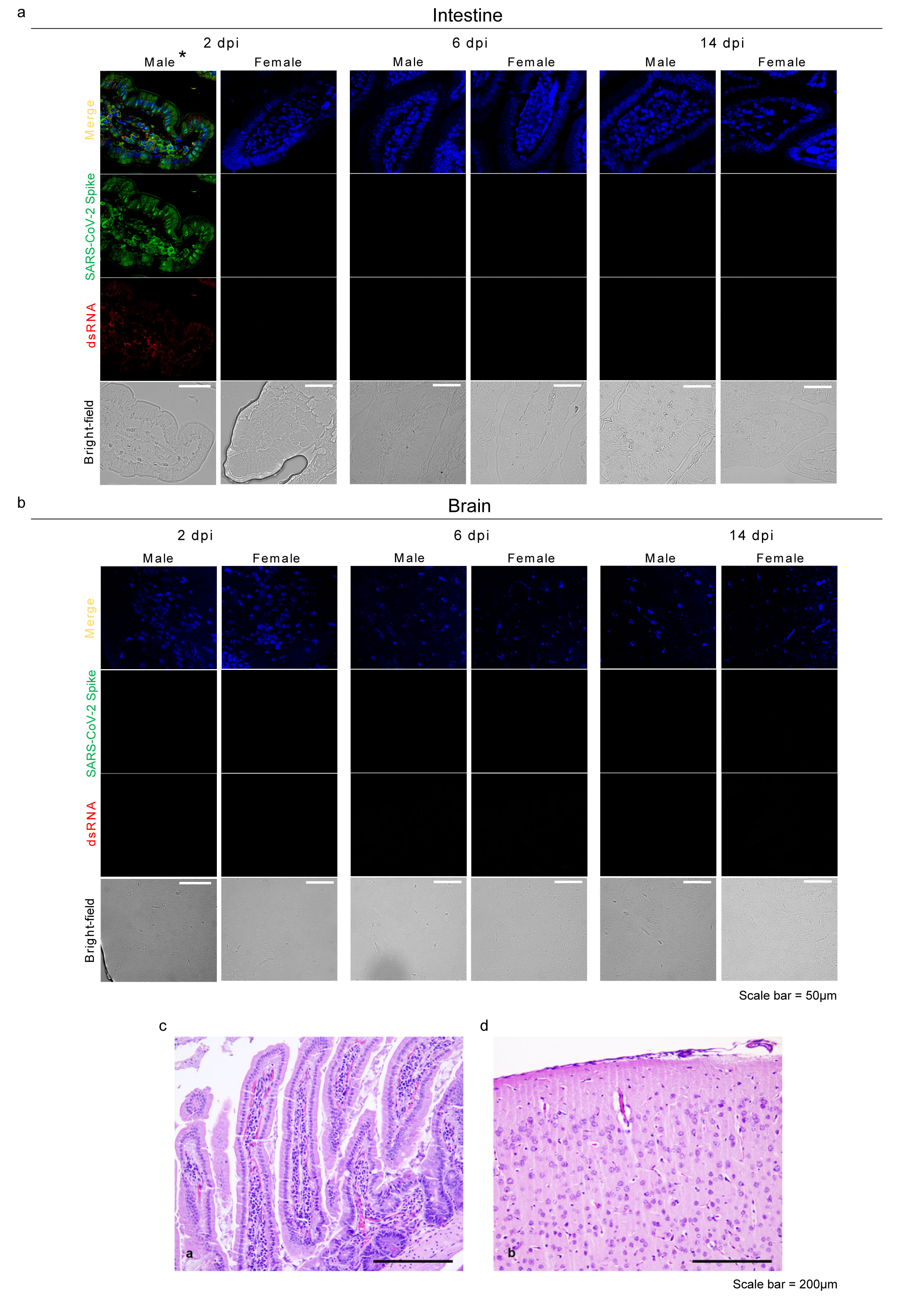

**Figure S1.** Representative immunofluorescence staining for SARS-CoV-2 Spike glycoprotein (green) and dsRNA (red) in infected male and female intestines (**a**) and brains (**b**). Scale bar = 50μm. All animals were analyzed, representative images are shown. **c-d**, Representative images of intestines (c**;** infected male 14 dpi) and brains (d**;** infected female 6 dpi) with no histopathological lesions. Scale bar = 200 μm. All animals were analyzed, representative images are shown. ***** is a representative image of one of the 2 out of 5 positive male intestines at 2 dpi; see the main text (Results section, Infection and seroconversion) for further details.

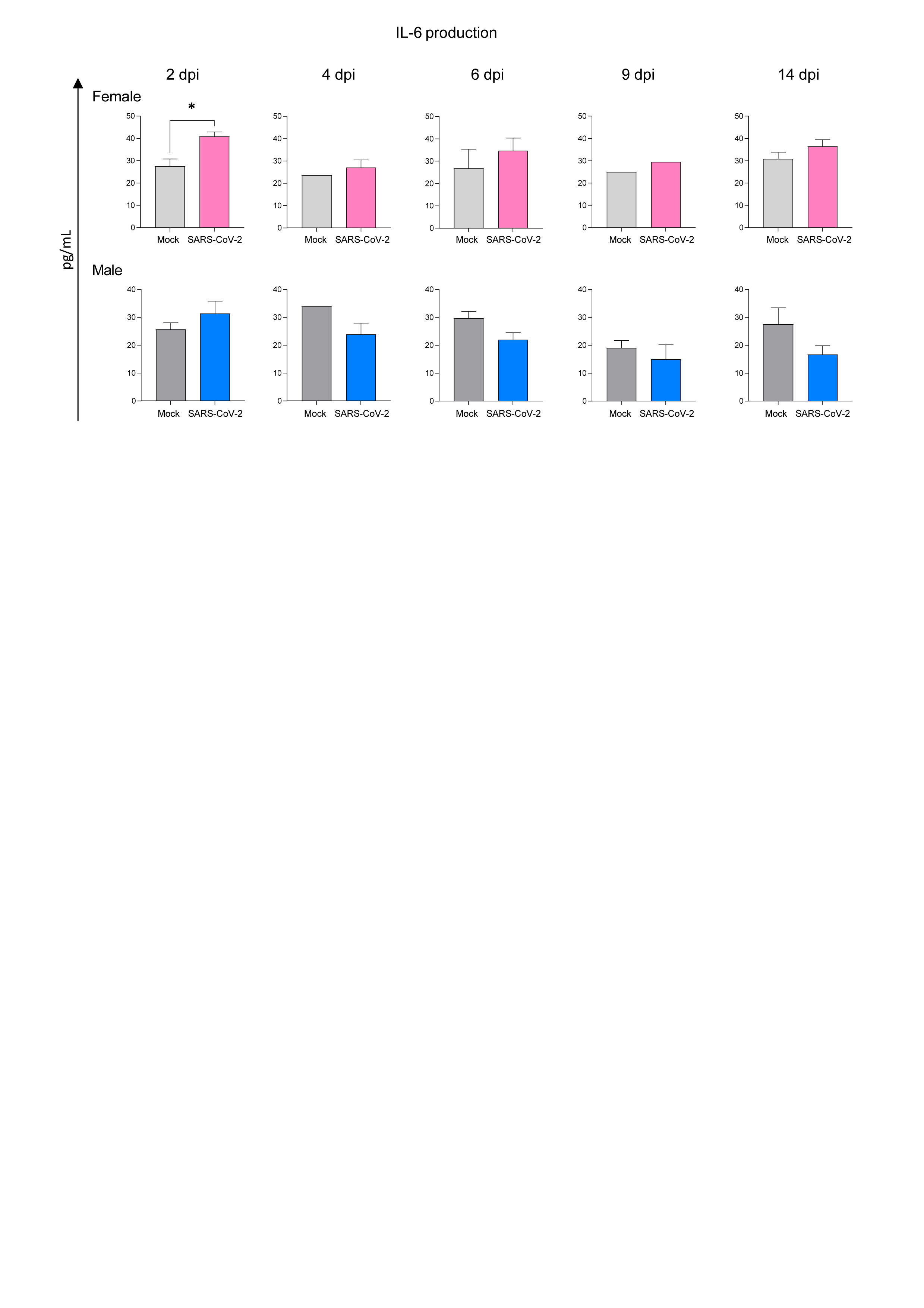

**Figure S2.** Singleplex ELISA levels (pg/mL) for mock and infected male and female hamsters IL-6. Wilcoxon-Mann-Whitney test of mock or SARS-CoV-2 M vs F; mean ± SEM is represented (* indicates a statistically significant comparison).

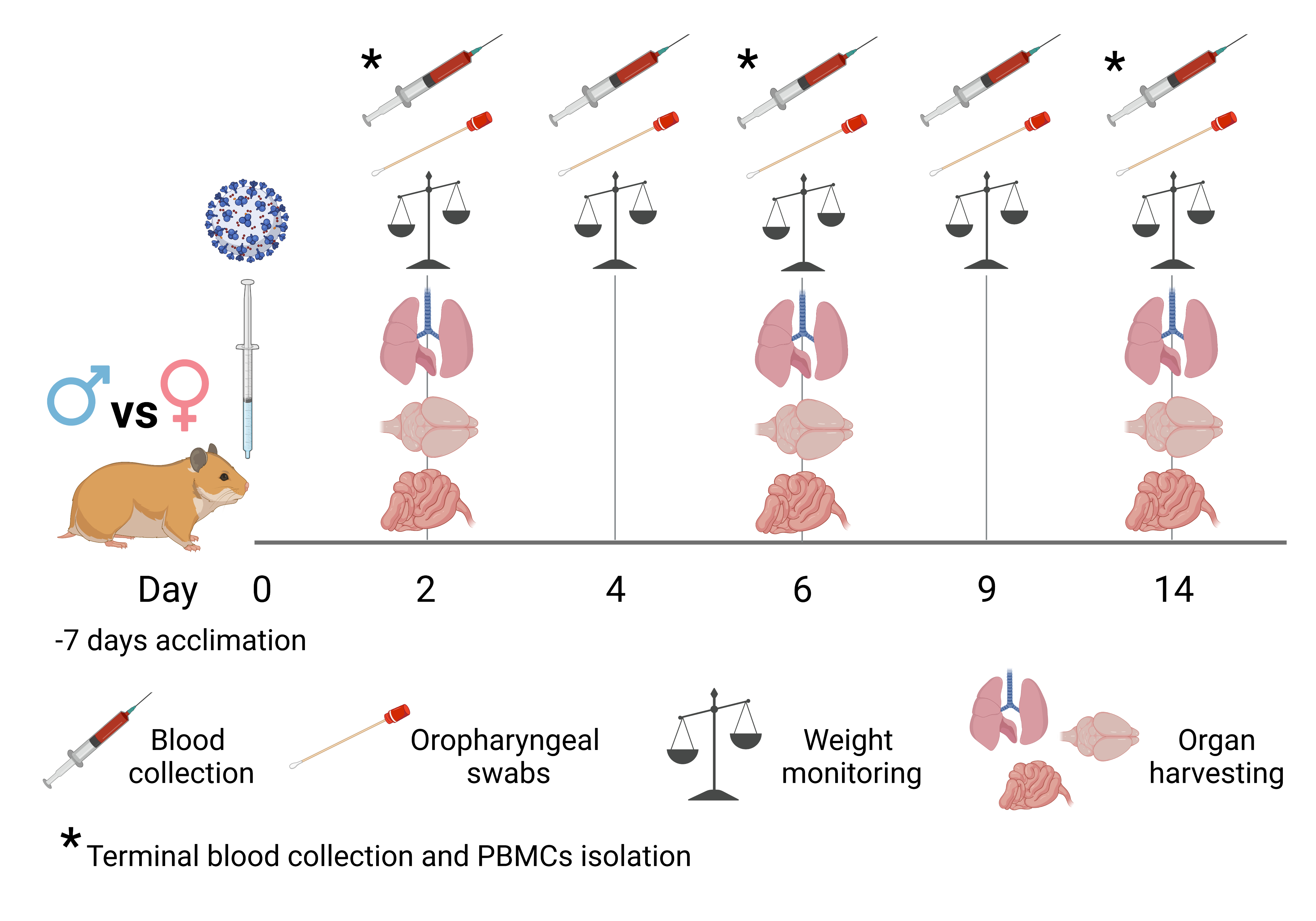

**Figure S3. The Syrian hamster as model for human SARS-CoV-2 infection.** *In vivo* study design. Created with BioRender.com (Agreement number HW24TOJLEV)

**Supplementary tables’ legends**

**Table S1.** Detailed statistical analysis.

**Table S2.** Histopathological scoring of H&E-stained lungs collected from male and female hamsters at 2, 6, and 14 dpi and from mock animals, related to Figure 2.

**Table S3.** Numbers of DEGs and Log2FC/FDR values found for each comparison made in differential expression analysis for the lungs, intestine and brains.

**Table S4.** Numbers of DEGs and GFOLD (reliable Log2FC) values found for each comparison made in differential expression analysis for PBMCs.

**Table S5.** Numbers of significant enriched GO terms and their scores found for each comparison made in differential expression analysis. GO terms are presented as blocks of the most specific term (i.e. the ones with the highest level) and their parent terms, based on the child-father relationships characterizing the GO graph. As a result, many GO terms, mostly the general ones, appear repeated in more than one block.
